# Dopamine modulates visual threat processing in the superior colliculus via D2 receptors

**DOI:** 10.1101/2021.02.12.430615

**Authors:** Quentin Montardy, Zheng Zhou, Lei Li, Qingning Yang, Zhuogui Lei, Xiaolong Feng, Shanping Chen, Qianqian Shi, Huiqi Zhang, Shuran Chen, Zhijian Zhang, Binghao Zhao, Fuqiang Xu, Zhonghua Lu, Liping Wang

## Abstract

Dopamine (DA) system is intriguing in the aspect that distinct, typically opposing physiological functions are mediated by D1 dopamine receptors (Drd1) and D2 dopamine receptors (Drd2). Both Drd1+ and Drd2+ neurons were identified in superior colliculus (SC), a visuomotor integration center known for its role in defensive behaviors to visual threats. We hypothesized that Drd1+ and Drd2+ neurons in the SC may play a role in promoting instinctive defensive responses.

Optogenetic activation of Drd2+ neurons, but not Drd1+ neurons, in the SC triggered strong defensive behaviors. Chemogenetic inhibition of SC Drd2+ neurons decreased looming-induced defensive behavior, suggesting involvement of SC Drd2+ neurons in defensive responses. To further confirm this functional role of Drd2 receptors, pretreatment with the Drd2+ agonist quinpirole in the SC impaired looming-evoked defensive responses, suggesting an essential role of Drd2 receptors in the regulation of innate defensive behavior. Inputs and outputs of SC Drd2+ neurons were investigated using viral tracing: SC Drd2+ neurons mainly receive moderate inputs from the Locus Coeruleus (LC), whilst we did not find any incoming projections from other dopaminergic structures. Our results suggest a sophisticated regulatory role of DA and its receptor system in innate defensive behavior.

## Introduction

Defensive behaviors are essential for survival, and requires detection and optimal behavioral selection at the sensorimotor level. Dopamine (DA) is a neurotransmitter synthetized in a limited set of brain structures, including the zona incerta (ZI), the ventral tegmental area (VTA) and the locus coeruleus (LC) (Björklund and Dunnett, 2007). It is involved in the learning and prediction of aversive events (Cohen et al., 2012; de Jong et al., 2019; Matsumoto et al., 2016), in sensorimotor control (Barrios et al., 2020; Frau et al., 2016; Pérez-Fernández et al., 2017) and in action selection (Howard et al., 2017; Kardamakis et al., 2015). There is growing evidence which indicate DA’s involvement in defensive behaviors (Barbano et al., 2020; Luo et al., 2018), notably that there is a high correlation between signal saliency and uncertainty when expecting an incoming aversive stimulation (Fiorillo, 2003; Jo et al., 2018). Extending this idea, dopamine is thought to have a dynamic effect on action and behavior selection at the earliest levels of sensory integration (Essig and Felsen, 2016; Hoyt et al., 2019; Kardamakis et al., 2015). The superior colliculus (SC), a subcortical structure receiving direct retinal afferents (Basso and May, 2017; Sparks, 1986), is known for its role in early sensorimotor integration (Ito and Feldheim, 2018). The SC is also thought to detect stereotypical salient visual information, such as snakes (Almeida et al., 2015), crawling in primates (Almeida et al., 2015; Isbell, 2011; Le et al., 2016; Maior et al., 2011), and collision or airborne predators in mice (Yilmaz and Meister, 2013), before relaying the information over a few synapses to core emotional centers such as the amygdala (Shang et al., 2015; Wei et al., 2015; Zhou et al., 2019). Thus, in recent years, several pathways originating from the SC have been identified, revealing an SC-Pulvinar-Amygdala pathway controlling defensive behaviors (Wei et al., 2015), and an SC-VTA-Amygdala pathway controlling flight behaviors (Zhou et al., 2019). Additionally, SC dysfunction in the early detection of visual threats is thought to negatively contribute to emotional and psychiatric disorders, in particular to Post Traumatic Stress Disorder (PTSD) (Lanius et al., 2017; Nicholson et al., 2017; Rabellino et al., 2016).

Interestingly, expression of dopaminergic receptors in the SC have been reported in many species including lamprey (Pérez-Fernández et al., 2014), rodents (Bolton et al., 2015; Mengod et al., 1992), non-human primates (Ciliax et al., 2000) and humans (Hurd et al., 2001; Mengod et al., 1992). In mice, SC DA receptors are mainly Drd1 and Drd2 (Bolton et al., 2015), but their upstream targets remain elusive, and their function largely unknown. We hypothesized that SC neurons expressing dopaminergic receptors may be involved in defensive behaviors in response to visual threats.

## Results

### Optogenetic activation of Drd2+ neurons in the SC, but not D1R, induces immediate flight behavior

To determine whether Drd1+ and Drd2+ SC neurons are involved in the control of defensive-like behaviors, we used an optogenetic strategy. First, we unilaterally injected the Cre-dependent adeno-associated virus AAV-DIO-ChR2-mCherry into the SC of Drd1-cre and Drd2-cre mice expressing Cre recombinase, selectively targeting SC neurons expressing dopamine receptors D1 or D2. Following virus injection, an optical fiber was placed above the SC (Fig. 1.A, up). Analysis of virus expression revealed that Drd2+ neurons were mostly localized in the intermediate SC layers (Fig. 1.B), whilst Drd1+ neurons were mainly found in the superficial SC layers (Fig. 1.C), confirming that these two categories of SC neurons are mainly segregated by different layers. To understand the function of each type in the context of defensive behaviors, mice were placed in an open field with a nest as a hiding place. They were allowed to explore the apparatus for 3 min (Fig. 1.A, down) during a pre-stimulation period in which both D2-cre and D1-cre animals showed typical exploratory behavior (Fig. 1.D, left). Optogenetic stimulation was then delivered (2.5 s, 20 Hz), during which time D1-cre mice maintained normal activity yet D2-cre mice immediately fled to their nest before freezing inside for at least 30 s post-stimulation, (Fig. 1.D, supplementary video 1-2), an effect observed in every individual in the D2-cre group. Consist with this, only the D2::ChR2 group rapidly increased speed immediately following stimulation (Fig. 1.E). On average, when all groups were compared, only the D2::ChR2 mice had flight-to-nest behavior (latency: D1::ChR2: 22.97 ± 6.8 s; D2::ChR2: 0.59 ± 0.12 s; D2::mCherry 22.77 ± 3.92 s;**P=0.0052, **P=0.0109), shown by the latency to reach the nest after stimulation (Fig. 1.F). In addition, the average time spent in the nest after stimulation was similarly low for D1-cre and control D2-mCherry (D1::ChR2: 33.81 ± 10.31 %; D2::mCherry 27.89 ± 2.92 %;****P<0.0001), and was significantly higher for D2:: ChR2 mice (D2::ChR2: 99.01 ± 0.12 %;****P<0.0001). These data suggest that Drd2+, but not Drd1+, SC neurons can induce defensive behaviors. Supporting this idea, SC Drd2+ neuronal projections (Sup. Fig. 1.A-B) encompass structures such as the lateral pulvinar, the ventral tegmental area, the parabigeminal nucleus, and the periaqueductal gray. In summary, these results indicate the Drd2+ neurons are sufficient to trigger defensive behaviors.

**Figure 1.**
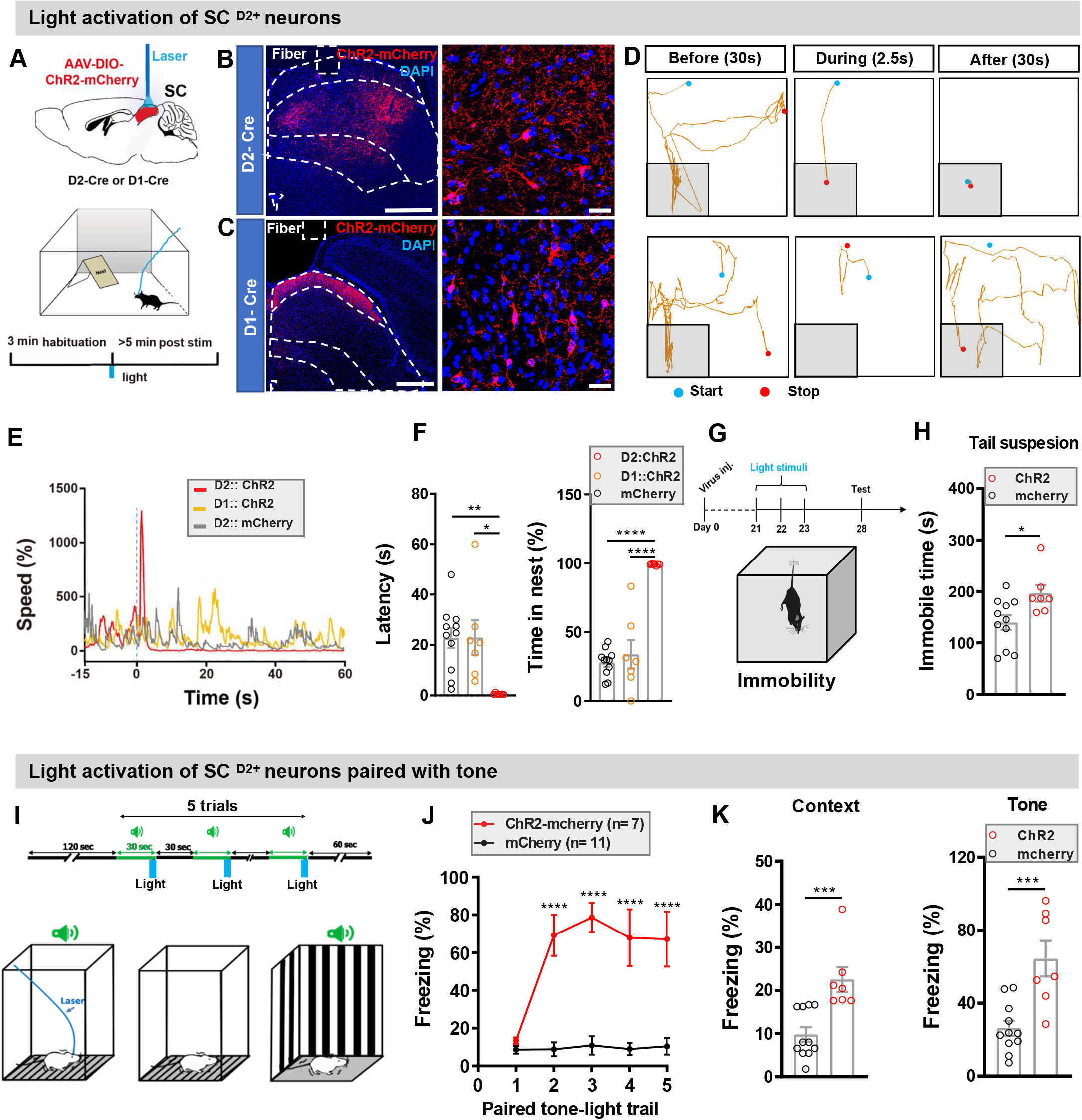
Optostimulation of SC ^D2+^ neurons induced strong defensive behaviors and fear memory. **(A)** Optogenetic strategy showing unilateral SC optical activation and experimental timeline. **(B-C)** Representative IHC shows selective targeting of ChR2-mCherry to SC ^D2+^ neurons (B) and SC ^D1+^ neurons (C), and the position of the fiber track (blue, DAPI; red, ChR2-mCherry; scale bars, 500 μm and 50 μm, respectively; solid line, fiber track). (**D**) Representative track plots of the SC ^D2+^ activated (up) and SC ^D1+^ (bottom) activated mice in open field with a nest demonstrating flight-to-nest defensive behavior of SC ^D2+^ activated mouse. (**E**) Representative speed profiles illustrate shorter flight latency after SC-D2+ activation in the ChR2-mCherry group than in the mCherry control group. (**F**) Following photostimulation of SC ^D2+^ neurons, the D2:: ChR2 group had lower flight latencies and higher time in the nest compared with controls (n _D2-mCherry_ =11 mice; n _D1-ChR2_ =7 mice, n _D2-ChR2_=7 mice; \*\**P* _*latency*_= 0.0032, *F*_*2, 22 latency*_=7.553, \*\*\*\**P* _*tim*e_< 0.0001, *F*_*2, 22 time*_= 48.9; Bonferroni *post hoc* test, for latency: D2-ChR2 VS. D2-mCherry, \*\**P* _*latency*_= 0.0052; D1-ChR2 VS. D2-ChR2, \*\**P* _*latency*_= 0.0109; for time in nest: D2-ChR2 VS. D2-mCherry, \*\*\*\**P* _*tim*e_< 0.0001; D1-ChR2 VS. D2-ChR2, \*\*\*\**P* _*tim*e_< 0.0001; one-way ANOVA). For all graphs, data were presented as mean ± SEM. (**G**) Experimental procedure for the repeated activation of SC ^D2+^ neurons caused depression-like behavior as indexed by elevated freezing. (**H**) Repeated activation of SC ^D2+^ neurons induced significant higher immobility time in the ChR2 group than in the control group (n _mCherry_ = 11 mice, n _ChR2_ =7 mice, *t* _*16*_= 2.569, \**P=*0.0206; Unpaired student test). (**I**) Schematic of the conditioned paring of activation of SC ^D2+^ neuronal activation and the tone. (**J**) Optogenetic stimulation SC ^D2+^ neurons increased freezing levels during conditioning (n _D2-mCherry_ =11 mice; n _D2-ChR2_=7 mice; Group x trial effect interaction, *F*_*4, 19*_ =11.77, \*\*\*\**P* < 0.0001, two-way ANOVA bonferroni *post hoc* test, \*\*\*\**P* < 0.0001). (**K**) Testing day: compared with D2-mCherry group, the D2-ChR2 group had a significantly higher percentage of freezing time in context (K-left) and tone (K-right) memory retrieval (n _mCherry_ = 11 mice, n _ChR2_ =7 mice, for context, *t* _*16*_= 4.13, \*\*\**P=*0.0008; for tone, *t* _*16*_=4.132, \*\*\**P=*0.0018; unpaired student test).

### Repeated activation of SC Drd2+ neurons induces long-term memory and depression-like behavior

To determine whether SC-Drd2 stimulation induces simple behavioral patterns or long-term emotional states, we investigated whether aversive stimulation elicits long-term affective states. To do this, we first used repeated activation of SC Drd2+ neurons to understand if would lead to depression-like behavior. In detail, ChR2 and mCherry control groups received 2.5 s repeated optogenetic stimulation for 3 consecutive trials (20 Hz, 5 ms pulse duration, 5-8 mW, 1 min interstimulus interval) over 3 consecutive days (Fig. 1.G). Five days after the previous session, a tail-suspension test revealed that the ChR2 group remained immobile significantly longer than the mCherry control group (Fig. 1.H; ChR2: 196.1± 42.72 s, mCherry: 139.5± 47.21 s; *P=0.0206), confirming that Drd2+ neurons can trigger long-term emotional states.

We next investigated whether SC Drd2 neuronal stimulation could lead to the formation of long-term aversive memories. To answer this question, we placed mice in a contextualized box to undergo classical Pavlovian conditioning (Fig. 1.I). Mice received an 80 dB tone over 30 s conditioned stimulus (CS) terminated with a 2.5s 20 Hz optogenetic stimulation of Drd2+ SC neurons as an aversive unconditioned stimulus (US). Mice were placed in the same context without tone delivery 24 h later or placed in a different context with tone delivery. During tone presentation during the conditioning trial, freezing time for all animals was significantly higher in the ChR2 test group than in the mCherry control group (ChR2: 69.18% ± 10.91%, mCherry: 8.75% ± 3.68%; ****P< 0.0001), confirming that SC Drd2+ neurons activation promote defensive behaviors (Fig. 1.J). During context retrieval, the ChR2 group spent significantly longer freezing than the mCherry controls (ChR2: 22.55 ± 2.87%, mCherry: 9.81 ± 1.67 %; ***P=0.0008) (Fig. 1.K, left). Similarly, in a different context presentation of CS stimulation alone led to freezing time being significantly higher in the ChR2 group than in the mCherry group (ChR2: 66.40% ± 9.79%, mCherry: 26.22 ± 4.06%; ***P=0.0008) (Fig. 1.K, right), overall indicating SC Drd2+ neuronal stimulation is aversive and can be used as an effective US during memory formation.

Overall, these results suggest that Drd2+ neurons can not only trigger defensive responses, but are also sufficient to promote formation of conditioned memories, and provoke long-term depression-like behaviors.

### Chemogenetic inhibition of D2R neurons impairs defensive behavior to looming stimuli

To question whether Drd2+ SC neurons are necessary to process visually-induced instinctive defensive behaviors, we unilaterally injected AAV vectors containing the chemogenetic inhibitory hM4Di receptors (AAV-DIO-HM4Di-mCherry) in the SC of Drd2-cre mice (Fig 2.A). Robust expression of mCherry was observed in the intermediate layers of the SC (Fig. 2.B). Instinctive defensive behaviors were elicited by placing mice in a box with a hiding nest, and by presenting an overhead looming stimulation known to result in a rapid flight response (Yilmaz and Meister, 2013). One hour before stimulation, HM4Di-test and mCherry-control groups received an IP injection of clozapine-*N*-oxide (CNO) (Fig. 2.A). During looming stimulation, flight latency was significantly higher in the HMDi-test group than in the control group (Fig 2.C; HM4Di: 3.54 ± 0.56s; mCherry: 2.25 ± 0.22s; *P=0.034). There was a non-significant trend for mice in the HM4Di-test group to spend less time in the nest than those in the control group, and the percentage of flight following stimulation was similar between groups (Fig. 2.D). This indicates that inhibition of Drd2+ SC neurons disrupts defensive behaviors to visual threats.

**Figure 2.**
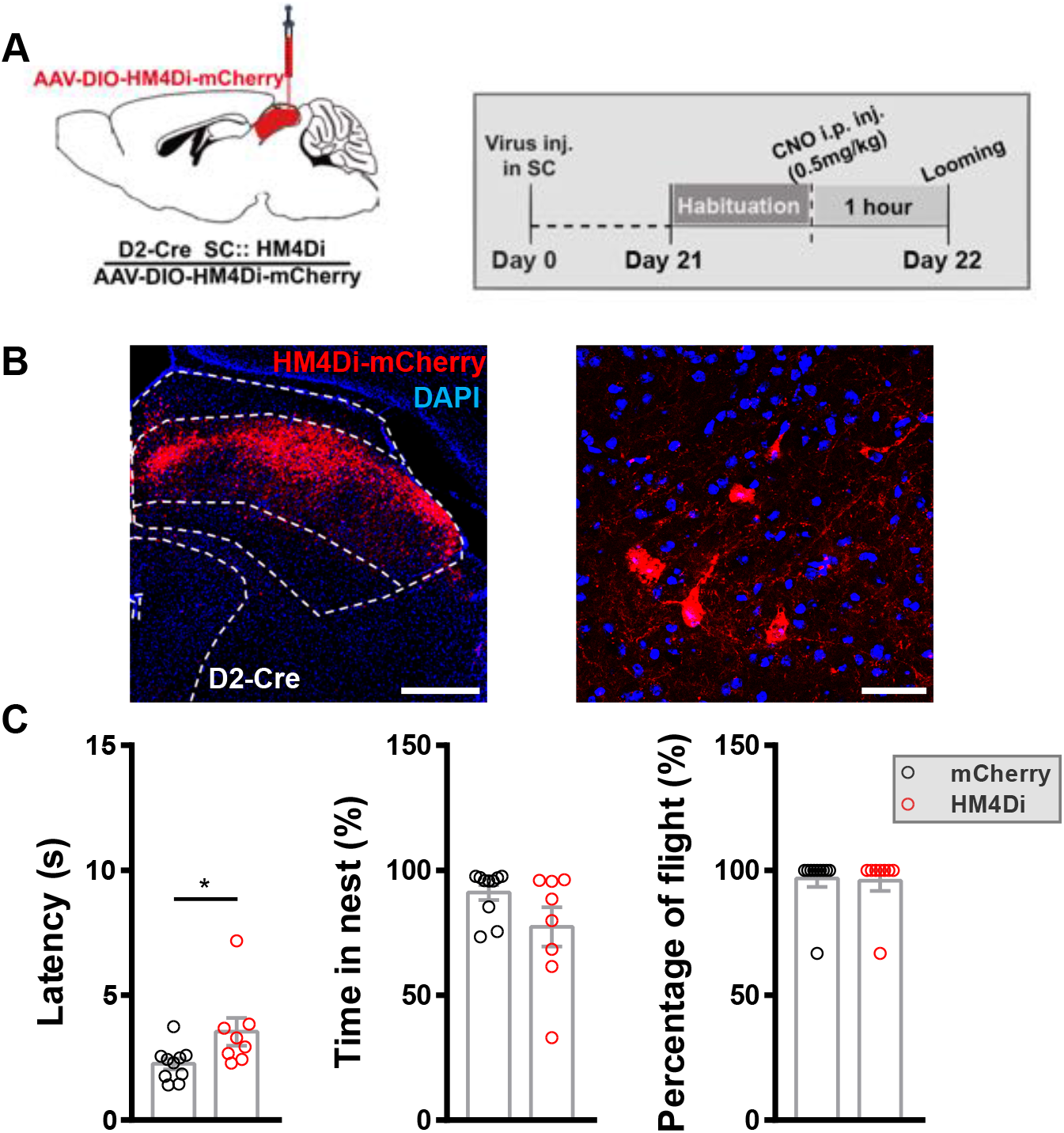
Chemogenetic inhibition of SC ^D2+^ neurons decreased looming-induced defensive behavior. (**A**) Chemogenetic strategy showing bilateral SC inhibition and experimental timeline. (**B**) Representative IHC showing selective targeting of hM4Di-mCherry to SC ^D2+^ neurons (blue, DAPI; red, hM4Di-mCherry; scale bars, 200 μm and 20 μm, respectively). (**C**) After CNO administration, the flight latency in the hM4Di group was higher than the mCherry controls (n _mCherry_ = 10 mice, n _HM4Di_ = 8 mice, for latency, *t* _*16*_= 2.326, \**P=*0.0335; for time in nest, *t* _*16*_=1.769, *P=*0.0959; for percentage of flight, *t*_*16*_=0.1582, *P=*0.8762; unpaired student t test). For all graphs, data are presented as mean ± SEM.

### Bilateral dopamine agonist quinpirole injection in the SC disrupts defensive responses to looming stimuli

We next wanted to investigate the net effect of dopamine in the superior colliculus in the context of defensive behaviors, and in particular, whether dopamine could modulate Drd2+ SC neuronal activity following visual threat. To do so, we first used patch-clamp slice recordings to characterize the effect of dopamine on Drd2 neurons. By injecting an AAV-DIO-EYFP virus into the SC of Drd2-cre mice, neurons were determined and patched on slice based on fluorescence (Fig. 3.A). Quinpirole, a selective D2 receptor agonist, was then delivered to the cells resulting in suppression of Drd2+ SC neuronal activity, with firing rate drastically reduced compared to baseline levels (100% VS. 14.54%) (Fig. 3.B). Next, to understand the physiological role of dopamine on defensive behaviors to visual threats, we bilaterally injected quinpirole or saline solution into the SC of wild type mice, and then presented looming stimulation 30 minutes later (Fig. 3.C). Flight latency was significantly shorter (Quinpirole: 14.7 ± 5.0 s ; Saline: 2.4 ± 0.3s; *P=0.026) and the probability for mice to flight to nest (Quinpirole: 64.2± 9.6%; Saline: 95.3 ± 3.2%; **P=0.0049) was significantly longer in the quinpirole group than in the saline control group, whilst time in the nest remained similar (Fig. 3.D). This confirms that dopamine modulates the SC activity and decreases defensive responses to aerial visual threat.

**Figure 3.**
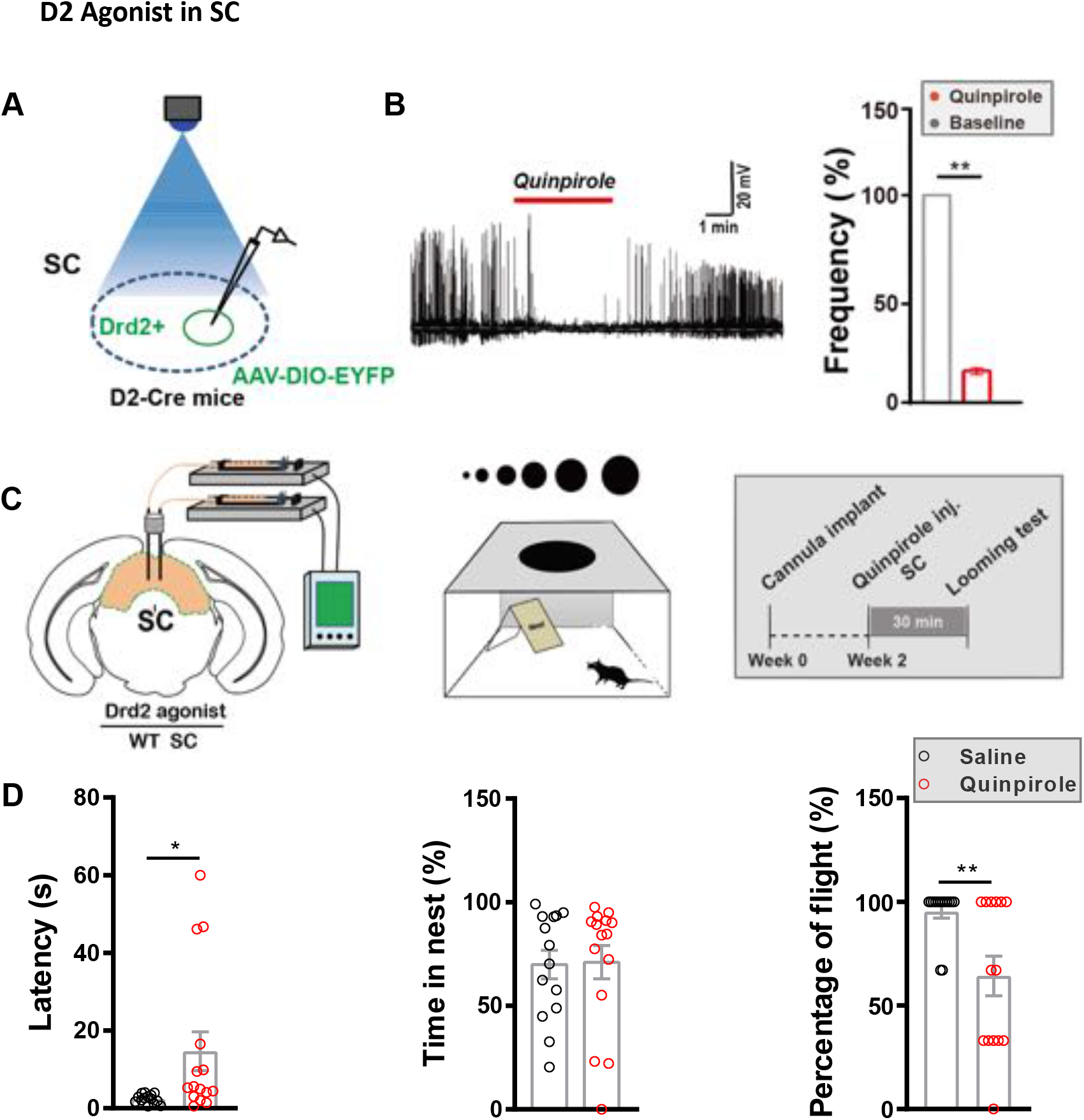
Drd2 agonist suppressed SC ^D2+^ neurons firing at brain slice recording intra-SC, and SC infusion dampened the looming-induced defensive behaviors in vivo. **(A)** Schematic showing in vitro patch-clamp slice recording of single-unit SC-D2+ neuronal activity following Drd2 agonist injection into the SC. AAV-DIO-EYFP injections in D2-cre mice were used to visualize D2-positive neurons. **(B)** *Right*, representative example of firing rate showing that the activity of SC-D2+ neurons was suppressed after infusion with Drd2 agonist; *left*, quantification of the firing rate of SC-D2+ neurons (n= 3 cells from 3 mice, data presented as mean ± SEM, **P=0.0033, t 2= 17.30, Paired student t test) **(C)** Bilateral Drd2 agonist strategy showing bilateral SC agonist infusion and experimental timeline. **(D)** The looming-induced flight-to-nest behavior was reduced by intra-SC infusion of Quinpirole (dopamine receptor 2 agonist), resulting in a recovery of flight latency and lower percentage of flight-to-nest (n _saline_ = 14 mice, n _Quinpirole_ = 14 mice, for latency, *t* _*27*_= 2.353, \**P=*0.0262; for time in nest, *t* _*27*_=2.372, *P=*0.909; for percentage of flight, *t*_*27*_=3.007, \*\**P=*0.0049; Unpaired student t test). For all graphs, data are presented as mean± SEM.

These blunted behavioral responses to visual threats suggest that SC D2 receptors are involved in triggering instinctive defensive behaviors to visual threats, and are necessary for the normal expression of the full repertoire of mouse behavior.

### The LC is the principal candidate sending dopaminergic projections to SC Drd2+ neurons

Finally, to determine the source of dopaminergic modulation in the SC, we injected the retrograde tracer cholera toxin B (CTB) into the SC (Sup. Fig. 2.A). CTB tracer was found in structures such as the primary visual cortex and the anterior cingulate cortex, both know to project to the SC (Baldwin et al., 2019; Zingg et al., 2017) (Sup. Fig. 2.B1-B2). CTB tracer was also found in the ventral tegmental area (VTA), the substantia nigra (SN), the periaqueductal gray (PAG), the paraventricular nucleus of the hypothalamus (PVN), or the dorsal raphe nucleus (DRN), known to synthetize dopamine (Björklund and Dunnett, 2007) (Sup. Fig. 2.C1-C5, D1). Finally, we found strong CTB fluorescence retrograde tracer signal in neurons in the locus coeruleus (LC), as well as the zona incerta (ZI) (Sup. Fig. 2.D2-D3). Together, these data demonstrate that SC receives numerous projections from dopaminergic structures, confirming previous reports which used equivalent methods to show that ZI and LC to be a source of dopamine in SC (Bolton et al., 2015). But retrograde tracer injection of CTB is not specific to dopaminergic projections to SC Drd2 neurons. To determine the dopamine source of the SC neurons expressing dopamine receptor D2, which possibly modulates defensive behaviors to visual threat, we mapped projections upstream from Drd2+ SC neurons using a Cre-dependent monosynaptic retrograde tracing technique. Drd2 -Cre transgenic mice received AAV-CAG-DIO -TVA-GFP (AAV2/9) and AAV-CAG-DIO-RG (AAV2/9) virus injections into SC. Three weeks after virus injection, the SC was infected with RV-EvnA-DsRed (EnvA-pseudotyped, G-deleted and DsRed-expressing rabies virus) using the same coordinates (Fig. 4.A). Whole brains were sectioned and stained with the fluorescent dopamine synthesizing enzyme tyrosine hydroxylase (TH) to confirm upstream neurons were capable of dopamine production. We found that the TVA-GFP and RV viruses were expressed in the intermediate layers of SC (Fig. 4.B). Neurons co-expressing RV retrograde virus and TH immunofluorescence were found in the locus coeruleus of every mouse (Fig. 4.C) with 78.34 ± 9.72 % retrogradely labeled neuron being TH positive (Fig. 4.J), indicating that the LC sends dopaminergic projections to Drd2 SC neurons. Neurons in other dopaminergic structures such as the DRN, ZI, VTA, SN, PAG, or the arcuate nucleus also retrogradely expressed RV but did not co-express TH fluorescence (Sup. Fig. 3.D-G We did not find dopaminergic inputs to SC Drd2+ neurons using this method (Sup. Fig. 3.J). Together, these results suggest dopamine projections to Drd2+ SC neurons could mainly come from LC.

**Figure 4.**
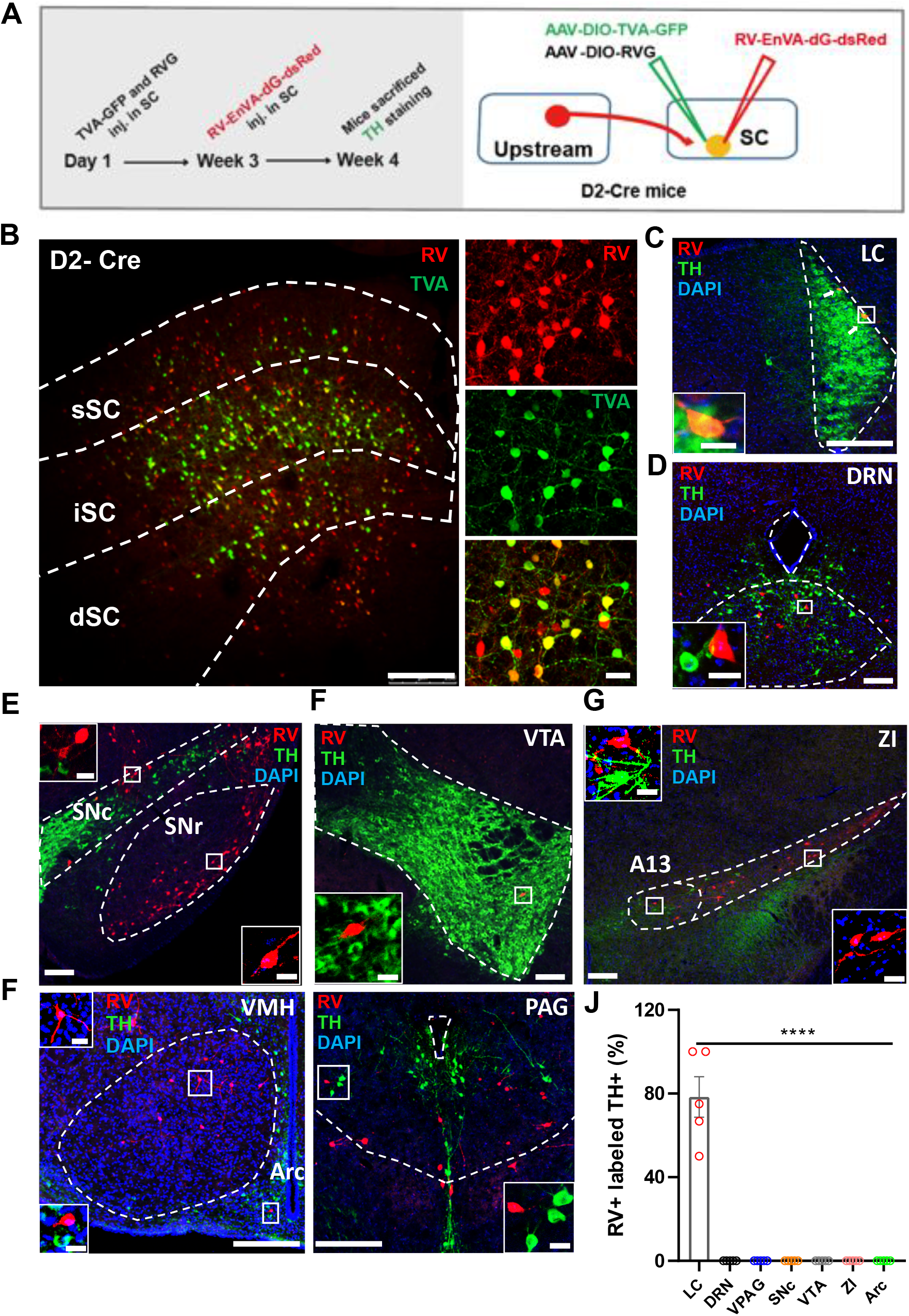
SC ^D2+^ neurons receive direct monosynaptic TH-positive inputs from LC. **(A)** Schematic of the rabies virus-based cell-type-specific monosynaptic tracing protocol. **(B)** Representative images denoting the starter cells in the SC of D2-Cre mice (Red, rabies-dsRed; green, TVA; blue, DAPI; scale bar, 250 μm and 25 μm, respectively). **(C-F)** SC-D2 RV retrograde labelled upstream brain regions and co-labelling with TH. Retrograde labelled cells (Red) in the (C) Locus coeruleus (LC), (D) Dorsal raphe (DRN), (E) Substantia nigra, compact part(SNc), Substantia nigra, reticular part(SNr), (F) Ventral tegmental area (VTA), (G) Zona incerta (ZI), (H)Ventromedial hypothalamic nucleus(VMH), Arcuate hypothalamic nucleus (Arc) and (I) Periaqueductal gray(PAG) with inputs to SC-D2+ neurons, (Red, rabies-dsRed; green, TH; blue, DAPI, scale bar, 250 μm and 20 μm respectively). **(J)** Quantification of the percentage of rabies-dsRed labeled neurons that overlap with TH in regions upstream of SC-D2+ cells. (n = 5 mice, *F* _*6, 28*_ = 65.01, \*\*\**P*< 0.0001, data presented as mean ± SEM; one-way ANOVA).

## Conclusion and Discussion

### Conclusion

We investigated the function of SC neurons expressing either Drd2 or Drd1dopaminergic receptors. Using optogenetic tools, we demonstrated that SC Drd2, but not Drd1, neuronal activation was able to induce strong defensive behaviors in the absence of threatening stimuli, and long-term effects such as fear memory and depression-like behaviors. Both chemogenetic inhibition using the HM4Di-CNO system, and physiological inhibition using the D2 receptor agonist quinpirole in vivo, impaired defensive behaviors to visual threats. Interestingly, CTB retrograde tracers revealed that SC receives projections from dopaminergic brain structures, results then extended by RV tracing showing that SC Drd2+ neurons receive transsynaptic dopaminergic projections from LC TH^+^ neurons. These results suggest an essential and sophisticated role of dopamine in the SC, and more specifically, of the dopamine -Drd2 receptor system in regulating instinctive defensive behaviors

## Discussion

### SC Drd2 function

Whilst dopamine D1 and D2 receptor expression in the mice superior colliculus has been reported, their function was largely unknown. Here, we revealed that optogenetic activation of SC Drd2+ neurons induced short and long-term defensive behaviors. These results are in line with previous reports showing that direct SC stimulation can induce fear-like behavior in many species (Shang et al., 2015; Wei et al., 2015; Zhou et al., 2019). Supporting this idea, we have shown that Drd2+ neurons are mainly localized in the intermediate layers of the SC (Fig. 1.B), the same SC layers that project to downstream structures involved in defensive and promoting flight behavior (Evans et al., 2018; Zhou et al., 2019). Indeed, AAV tracing of SC Drd2 neurons demonstrated a projection to structures known to receive SC inputs that control defensive behaviors (Zhou et al., 2019). Drd2+ neurons, of which a major proportion are excitatory, enrich the intermediate layer of the SC, (May, 2006) (Bolton et al., 2015). It is therefore possible that optogenetic activation of SC Drd2 neurons activates downstream nuclei involved in defensive behaviors, directly driving flight behaviors. In line with presented evidence showing that Drd2 neuronal stimulation is able to trigger flight in the absence of visual threat, inhibition of these neurons impairs defensive behavior following visual threat. Indeed, we demonstrated that quinpirole injection into the SC weakens defensive responses, in particular by increasing flight latency and decreasing the flight probability. However, defensive behavior was not only diminished, indicating that disruption of SC Drd2 neurons alone is not sufficient to abolish defense. It is therefore likely that the SC Drd2 neurons do not encompass all of the SC neurons that project to emotion-related structures downstream.

In parallel, when exploring the function of Drd1 SC neurons, we found that this subpopulation is not involved in triggering defensive behaviors. Given that Drd1 and Drd2 receptors may have different on behavior (Liu et al., 2019; Smith and Kabelik, 2017; Tu et al., 2019; Verharen et al., 2019), it may have been expected that SC Drd1 neurons facilitate action to generate flight execution, but this not been observed here. Neurons with Drd1 and Drd2 receptors do not always participate in the same function (Smith and Kabelik, 2017). For example, D2, but not D1, neurons modulate auditory responses in the inferior colliculus (Hoyt et al., 2019). It is therefore reasonable to think that the function of Drd1 SC neurons may simply remain masked; indeed, dopamine at the SC level may have a broader scope of action than fear, such as the integration of visual signals among which looming-mimicking collisions are only a subset. Thus, to understand Drd1 SC role it would be necessary in the future to study the effects of dopamine on other non-emotional and more classical functions of the colliculus such as visuo-motor integration (Isa and Saito, 2001; Marino et al., 2008; Munoz et al., 1991) and attention selection (Ding et al., 2019; Evans et al., 2018; White et al., 2019).

Finally, we can suppose that a portion of the SC neurons expressing dopaminergic receptors act more locally at the microcircuitry level. Knowing that 37% of the Drd2 neurons are GABAergic (Bolton et al., 2015), it is likely that most form local inhibitory projections at the microcircuit level (May, 2006; Tardif et al., 2005; Villalobos et al., 2018; Vokoun et al., 2010). Such local interactions are necessary to control visual integration (Kasai and Isa, 2016; Muller et al., 2018; Vokoun et al., 2010) and have even been associated with visual attention (Hafed et al., 2009). Knowing then, that a proportion of these Drd2 neurons are GABAergic, and that we confirmed that DA inhibits Drd2 SC neurons, we can reasonably propose that DA can also act by removing local inhibition. Weakened defensive responses could therefore be partly due to a release of lateral inhibition in the SC (Kasai and Isa, 2016), disrupting the integration of the visual signal (or tuning it to optimize detection of specific spatio-temporal frequencies) and indirectly leading to a reduction of defensive responses. This raises the broader question of whether neuromodulation at the SC level disrupts visual perception or impairs subsequent selection of action.

It is essential now to explore the effect of dopamine on local SC micro-circuitry to determine whether it participates in visual signal integration, and which categories of behaviors it affects.

### On the circuitry aspect

Dopaminergic receptors at the SC level have been found in several species (Ciliax et al., 2000; Hurd et al., 2001; Mengod et al., 1992; Pérez-Fernández et al., 2014), suggesting that dopaminergic projections innervate the SC. In addition, it has been demonstrated using mice that dopaminergic projections from the ZI could target the SC (Bolton et al., 2015). In this study, the method used consisted of injecting latex microspheres, a retrograde tracer that has no particular affinity for neurons expressing dopaminergic receptors, into the SC. Thus, it demonstrated that ZI and LC could send DA projections to SC, but not that these projections target neurons expressing dopamine receptors. Here, we used an RV retrograde virus in conjunction with Drd2-cre mice, specifically mapping upstream pathways to SC neurons expressing DA receptors. We observed that several dopaminergic structures project towards Drd2 neurons, but we only found that the LC as sent dopaminergic projections to the SC (see Sup. Fig. 3.J). Since previous work revealed DA projections to SC, in particular from ZI, it will be necessary to carefully detail their connectivity patterns, and to understand their function.

Although we do not exclude the existence of other dopaminergic projections, we propose that the LC is the main source of dopamine for Drd2+ SC neurons, which is able to modulate defensive behavior. This hypothesis is in line with a previous report from our group demonstrating that the LC sends TH positive adrenergic projections to the SC (Li et al., 2018). But these projections modulated defensive behavior following physiological stress by increasing flight probability, whilst here, we show a decrease of fear-like behaviors. To explain this discrepancy, it is necessary to note that Lie et al. used a NE antagonist, whilst we used DA agonist. Extending this idea, and knowing that a majority of LC neurons are NE positive (Amaral and Sinnamon, 1977; Robertson et al., 2013), it has been demonstrated that LC terminals can co-release dopamine and adrenaline/noradrenaline (Devoto et al., 2005a, 2005b). Dissociating the effect of DA from the one of NE in the context of LC-SC projections is important, but represents a real technical challenge to date.

The LC-SC projections we revealed are only moderate and are unlikely to be responsible alone for the behavioral phenomena reported in the present article. In addition, our CTB data are in line with previous reports showing that the ZI sends dopaminergic projections to the SC. But these projections do not target SC neurons expressing dopaminergic receptors, raising the question of the mechanisms by which DA reaches dopaminergic receptors in the SC. Partly answering this question, it is known and discussed that dopamine does not necessarily follow canonical neurotransmission mechanisms, and could follow a volume transmission mode of delivery (Fuxe et al., 2015; Liu et al., 2018; Sulzer et al., 2016). First, in other brain structures it has been shown that dopamine can be highly localized at the extra-synaptic rather than synaptic level (Devoto et al., 2003), whilst dopamine receptors are sometimes located far from their release sites (Caillé et al., 1996). In addition, recent studies revealed that the secretion of dopamine does not only take place at the synapse level, but could take place en-passant along the dopaminergic axons (Liu et al., 2018). This supports the hypothesis that diffusion and dilution are the main modes of action of the transmitter (for review: (Cragg and Rice, 2004; Rice et al., 2011; Rice and Cragg, 2008), which could explain how DA reaches dopaminergic receptors in SC without necessarily directly targeting specific receptors. In parallel, it is important to note that only a few sets of dopaminergic boutons can effectively release DA, as is the case in the striatum where only a minority of DA vesicles can release the transmitter (Pereira et al., 2016). This suggest that dopaminergic pathways to SC, whether projecting to DA receptors or not, are not necessarily active. Research on these particular DA transmission processes are still in their infancy, but this may explain in part why a proportion of the dopaminergic projections to SC do not directly synapse with neurons expressing D2 receptors. However, if such DA diffusion appears to be the prevalent mode of action in SC, it would make the role of DA in SC circuitry more challenging to elucidate. Understanding how, and in which context, dopamine is dynamically released in the SC is a key question for the near future.

It is of great importance for an animal to be able to predict the occurrence of a potential threat. A non-exclusive way to solve this problem is to optimize the detection of a threatening signal at the earliest stages of visual processing. We hypothesized DA could play such a role in the SC, and demonstrated Drd2 SC neurons were able to induce defensive responses even in the absence of visual threat, and were necessary for the normal expression of an optimal behavior. Our results suggest DA and its receptors regulate innate defensive behaviors in a sophisticated manner. Still, understanding in which condition DA is released in the SC is of high important, especially by which global and local mechanisms DA reaches its SC receptors, and understanding the dynamics involved. Understanding how defensive behaviors can be modulated from the earliest perceptive stage could help to find new therapeutic solutions to psychiatric pathologies, such as post-traumatic disorders.

## Materials & Methods

### EXPERIMENTAL MODEL AND SUBJECT DETAILS

#### Animals

All husbandry and experimental procedures in this study were approved by the Animal Care and Use Committees at the Shenzhen Institute of Advanced Technology (SIAT) or Wuhan Institute of Physics and Mathematics (WIPM), Chinese Academy of Sciences (CAS). Adult (6 to 8 weeks old) male C57BL/6J (Guangdong Medical Laboratory Animal Center, Guangzhou, China), Drd1-Cre (MMRRC_030989-UCD), and Drd2 -Cre (MMRRC_032108-UCD) mice were used in this study. Mice were housed at 22–25 °C on a circadian cycle of 12-hour light and 12-hour dark with ad-libitum access to food and water.

## METHOD DETAILS

### Viral vector preparation

For optogenetic experiments, the plasmids for AAV2/9 viruses encoding *EF1α*:: DIO-hChR2 (H134R)-mCherry, *EF1α*:: DIO-HM4Di-mCherry and *EF1α*:: DIO-mCherry were used. Viral vector titers were in the range of 3-6×10^12^ genome copies per ml (gc)/ml and viruses were all packaged by BrainVTA Co., Ltd., Wuhan. For rabies tracing, the viral vectors AAV2/9-*EF1α*:: DIO-TVA-GFP, AAV2/9-*EF1α*:: DIO-RV-G, and EnvA-RV-dG-dsRed were used and were all packaged by BrainVTA Co., Ltd., Wuhan. For retrograde tracing, AAV and rabies viruses were purified and concentrated to titers at approximately 3×10^12^ v.g /ml and 1×10^9^ pfu/ml, respectively.

### Virus injection

Mice were placed in a stereotaxic apparatus (RWD, China) before being anesthetized with pentobarbital (i.p., 80 mg/kg). Anesthesia was then maintained with isoflurane (1%) during surgery and virus injections. Injections were conducted with a 10 μl syringe (Neuros; Hamilton, Reno, USA), using a microsyringe pump (UMP3/Micro4, USA). Coordinates for virus injection of the SC in Drd2-Cre mice (total volume of 350 nl) were: bregma -3.80 mm, lateral ± 0.80 mm and dura -1.80 mm. SC in Drd1-Cre mice (total volume of 200 nl) coordinates were: AP -3.40 mm, ML ±0. 50 mm, and DV -1.5 mm. Viruses were delivered unilaterally for ChR2 and bilaterally for HM4Di.

### Trans-synaptic tracer labeling

All animal procedures were performed in Biosafety level 2 (BSL2) animal facilities. To determine whether the inputs of Drdr2+ and Drd1+ neurons in the SC, Drd2-Cre mice and Drd1 were used for trans-mono-synaptic tracing based on the modified rabies virus. A mixture of AAV2/9-*EF1α*:: DIO-RV-G and AAV2/9-*EF1α*:: DIO-TVA-GFP (1:1, total volume of 200-250 nl) was injected into the SC region. For virus injection into the SC in Drd2-Cre mice (total volume of 250 nl), the following coordinates were used: AP -3.80 mm, ML ± 0.80 mm and DV -1.80 mm. Coordinates for SC injections in Drd1-Cre mice (total volume of 200 nl) were: AP -3.40 mm, ML ±0. 50 mm, and DV -1.5 mm. Three weeks later, 200 nl of EnvA-RV-dG-dsRed virus was injected into the same coordinates in these mice. Mice were sacrificed one week after RV injection.

### Implantation of optical fiber(s) and cannulas

A 200 µm optic fiber (NA: 0.37; NEWDOON, Hangzhou) was unilaterally implanted into the SC in Drd2 mice (AP, -3.8 mm; ML, -0.6 mm; DV, -1.4 mm) and SC in Drd1 mice (AP, -3.40 mm; ML, -0.5 mm; DV, -1.0 mm). For pharmacological experiments, drug cannulas were bilaterally implanted into the SC (AP, -3.8 mm; ML, ±0.6 mm; DV, -1.4 mm). The mice were used for behavioral tests at least 1-2 weeks after surgery.

### Patch-clamp electrophysiology

Coronal slices (300 μm) containing the SC were prepared, using standard procedures, from 14-16 week-old Drd2-Cre mice, which had received virus injections three weeks earlier. Recordings in SC Drd2+ cells were made on visually identified neurons expressing EYFP.

Brain slice were cut using a vibratome (Leica) into a chilled slicing solution (in mM: 1.3 NaH_2_PO_4_, 25 NaHCO_3_, 110 Choline Chloride, 0.6 Na-Pyruvate, 0.5 CaCl_2_, 7 MgCl_2,_ 2.5 KCl, 1.3 Na-Ascorbate). Then, slices were incubated at 32 °C for 30 min in artificial cerebrospinal fluid (ACSF) (in mM: 10 Glucose, 2 CaCl_2_, 1.3 MgCl_2,_ 125 NaCl, 2.5 KCl, 1.3 NaH_2_PO_4_, 25 NaHCO_3_, 1.3 Na-Ascorbate, 0.6 Na-Pyruvate, pH 7.35) and allowed to equilibrate to room temperature for >30 min. The osmolarity of all solutions was maintained at 280–300 mOsm.

For current clamp, pipettes were filled with a solution (in Mm: 105 Cs-gluconate, 10 phosphocreatine (Na), 0.07 CaCl2, 4 EGTA, 10 HEPES, 4 Na-ATP, 1 Na-GTP, and 3 MgCl2). To identify the spike dopaminergic nature, D2 agonist quinpirole (10 μM) was added at the end of recordings.

Pipettes with a resistance of 3−5 MΩ were formed by a micropipette puller (Sutter P-2000). We viewed neurons with an upright fixed-stage microscope (FN-S2N; Nikon., Japan) during whole-cell patch recording with a MultiClamp700B amplifier (Molecular Devices). Analog signals were low-pass filtered at 2 kHz, digitized at 20 kHz using Digidata 1440A, and recorded using pClamp 10 software (Molecular Devices).

### Histology, immunohistochemistry, and microscopy

Mice were anesthetized with an overdose of chloral hydrate (10% W/V, 300 mg/kg body weight, i.p.) and were then transcardially perfused with PBS, followed by ice-cold 4% paraformaldehyde (PFA; Sigma) in PBS. Brains were extracted and submerged in 4% PFA at 4 °C overnight to post-fix. After pots-fixing, brains were transferred to 30% sucrose to equilibrate. Coronal slices (40 µm) were using a cryostat microtome (Lecia CM1950, Germany). Freely floating sections were incubated with PBS, containing blocking solution (0.3% TritonX-100 and 10% normal goat serum, NGS in PBS, 1 h at room temperature). Primary antibody (rabbit anti-TH, 1:500, Abcam) were incubated the slices. The antibody was diluted in PBS with 3% NGS and 0.1% TritonX-100 overnight. The secondary antibody Alexa fluor 488 (1:200, Jackson) was used to incubated at room temperature for 1 h. Slices were mounted and covered slipped with anti-fade reagent with DAPI (ProLong Gold Antifade Reagent with DAPI, life technologies) or signal enhancer (Image-iT FX Signal Enhancer, Invitrogen). All images were photographed and analyzed with a Leica TCS SP5 laser scanning confocal microscope and ImageJ, Image Pro-plus software.

For the rabies monsynaptic tracing, imagines were taken and then overlaid with The Mouse Brain in Stereotaxic Coordinates to locate the brain slices. Retrogradely identified positive neurons upstream of SC were manually counted by an individual experimenter blind to the experiment groups.

### Optogenetic manipulation

Before optogenetic stimulation, animals were handled and habituated for 10-15 min to the looming box with a nest shelter in corner one day before testing. During the test session, mice were put into the same looming box and allowed to freely explore the box for 3-5 min, then received 2.5 s of 473-nm blue laser (Aurora-220-473, NEWDOON, Hangzhou) with light power at the fiber tips (20 Hz, 5 ms pulse duration, 5-8 mW). Light stimulation was unilaterally delivered to the SC Drd1+ and Drd2+ cells without looming stimulation in this experiment. Light was presented twice at approximately 3-min intervals via a manual trigger. We manually triggered stimulation when mice were at the far end of the open field, away from the nest position, within one body-length distance from the wall.

### Fear conditioning

Fear conditioning was done over two sessions: a training session and a memory test. During the training session, mice were put into a fear conditioning chamber, located inside a sound-isolation box. The Drd2-cre mice were allowed to freely explore the chamber for 3 mins before an 85-dB, 2-kHz tone was presented for 30 s as conditioned stimulation (CS). This co-terminated with 2.5 secs light stimulations (20 Hz, 5 ms pulse duration, 5-8 mW) separated by 1-min intervals. Mice were kept in the training chamber for another 60 s before being moved outside. Each mouse received 5 repeated CS paired light stimulations.

During the memory test, mice performed consecutive tests: context test and tone test. 1) context test: mice were placed back into the altered chamber (modified by changing the white silver side walls to plastic walls decorated with black and white stripes, and changing the metal grid floor to a plastic sheet) for 5 mins to measure levels of freezing. 2) Tone test: an 80-dB, 2-kHz tone was presented for 1 min after the context test to measure freezing levels during the tone. 20% ethanol was used to clean the chamber to eliminate odors from other mice. All behavior were recorded and scored by the FreezeFrame fear conditioning system (Lafayette Instrument). Behavioral analysis was done blind to treatment group.

### Tail suspension test

Before the tail suspension test, Drd2-cre mice received repeated 2.5 s blue-light stimulation (20 Hz, 5 ms pulse duration, 5-8 mW) in the SC with 1-min intervals (3 times repeated light stimulation) in one day. This photostimulation was conducted for 3 consecutive days and then tail suspension tests were performed 7 days after last light stimulation.

During the test session, the tail suspension test was done in a 50 x 50 x 30 cm box with an open front. Mice were individually suspended by the tail with adhesive tape for 6 mins. An HD digital camera (Sony, Shanghai, China) positioned in front of the box was used to record behavior. Immobility were analyzed with Anymaze software (Stoelting Co.).

### Looming test and Pharmacological antagonism

A 40 x 40 x 30 cm closed Plexiglas box with a shelter nest in the corner was used for the overhead looming test. The looming box contained an LCD monitor on the ceiling to present a black disc expanding from a visual angle of 2° to 20° in 0.3 s, i.e., expanding speed of 60 °/s. Each looming stimulus included 15 repetitions of the expanding disc stimulus with a 0.066 s interval between each. Each looming stimulus lasted 5.5 s.

An HD digital camera (Sony, Shanghai, China) was used to record behavior. The behavioral test included two sessions, a pre-test and a test session. During the pre-test session, mice were handled and habituated for 10-15 min to the looming box one day before testing. During the test session, 200 nl saline (control) or D2 receptor agonists per hemisphere (Quinpirole, 0.25 μg /side) was bilaterally infused into the SC (AP, –3.8 mm; ML, ±0.6 mm; DV, –1.85 mm) 30 min before a looming test. Then, mice were put in the box and allowed to freely explore the box for 3-5 min. For pharmacological experiments plus looming, mice received 3 trials of looming stimulus but only defensive behavior to the first stimulus was analyzed; no observable adaptation was observed in any of the experiments.

### Behavioral analysis

Behavioral data were analyzed with Anymaze software. Individual time courses were plotted where T=0 ms as the time of stimulation. There three measures were obtained as indices of light-evoked or looming-evoked defensive behavior. (1) latency to return nest: the time from photostimulation or looming stimulus presentation to time when the mouse escaped/entered the nest; (2) time spent in nest (% of 1 min bin): time spent in the nest following looming stimulus or photostimulation; (3) percentage of flight (% of 3 repeated trial of looming stimulus). the probability of flight to nest after looming stimulus in 3 repeated photostimulation.

Flight is defined as episodes where speed increases 4 times than the average speed in cases where the final position is in the nest.

For all mice in this study, virus expression and fiber placements or cannula were confirmed by histological staining after our data were collected. Virus expression, behavioral tests and behavior analyses were performed by different experimenters. Decisions to discarded data on any given day was done blind to the behavioral groupings.

## QUANTIFICATION AND STATISTICAL ANALYSIS

All statistics were performed in Graph Pad Prism (GraphPad Software, Inc.), unless otherwise indicated. Paired student tests, unpaired student tests, and one-way ANOVAs were used and Bonferroni post hoc comparisons were conducted to detect significant main effects or interactions. In all statistical measures a P value <0.05 was considered statistically significant. Post hoc significance values were set as *P< 0.05, **P< 0.01, ***P< 0.001 and ****P< 0.0001; all statistical tests used are indicated in the figure legends.

## Acknowledgements

This work was supported by National Natural Science Foundation of China (NSFC) 31630031 (L.W .), NSFC 31930047(L.W .), NSFC 81425010 (L.W .), NSFC31971072 (L.L.); International Partnership Program of Chinese Academy of Sciences 172644KYS820170004 (L.W.); Helmholtz-CAS Joint Research Grant GJHZ1508 (L.W.); the Strategic Priority Research Program of Chinese Academy of Science, XDB32030200; Guangdong Provincial Key Laboratory of Brain Connectome and Behavior 2017B030301017 (L.W.); JCYJ20170413164535041(L.W.), JCYJ20150401150223647 (Z.Z.); Shenzhen Municipal Funding GJHZ20160229200136090 (L.W.); Shenzhen Discipline Construction Project for Neurobiology DRCSM [2016]1379 (L.W.);Science and Technology Planning Project of Guangdong Province 2018B030331001(L.W.); CAS President’s International Fellowship 2020FYB0005 (Q.M.); Guangdong Province International Scientific and Technological Cooperation 2019A050508008 (M.Q.).

## Author contributions

M.Q., Z.Z., and L.L. contributed equally to this work. M.Q., Z.Z., L.L. and L.W. designed and initiated the project. Z.Z. performed virus injections, fiber and cannula implantation. M.Q., Z.Z and L.L. setup the behavior protocol. Z.Z., X.F., Q.S., Z.L. (Zhuogui Lei), M.Q and L.L. performed behavior experiments. Z.Z., M.Q. and L.L. processed and analyzed behavior data. Z.Z. performed rabies virus injections. Q.Y., H.Z., S.C., and Z.Z. performed immunohistochemistry and quantitative analyzes of the tracing data. S.C. performed the patch clamp recording. Z.L. (Zhonghua Lu) provided the viral vectors. Q.M., Z.Z., L.L. and L.W. interpreted the results. M.Q., Z.Z and L.L. wrote the manuscript. L.W. supervised all aspects of the project.

## Supplementary Figure Legends

## Figure Legends

**Supplementary Figure 1.**
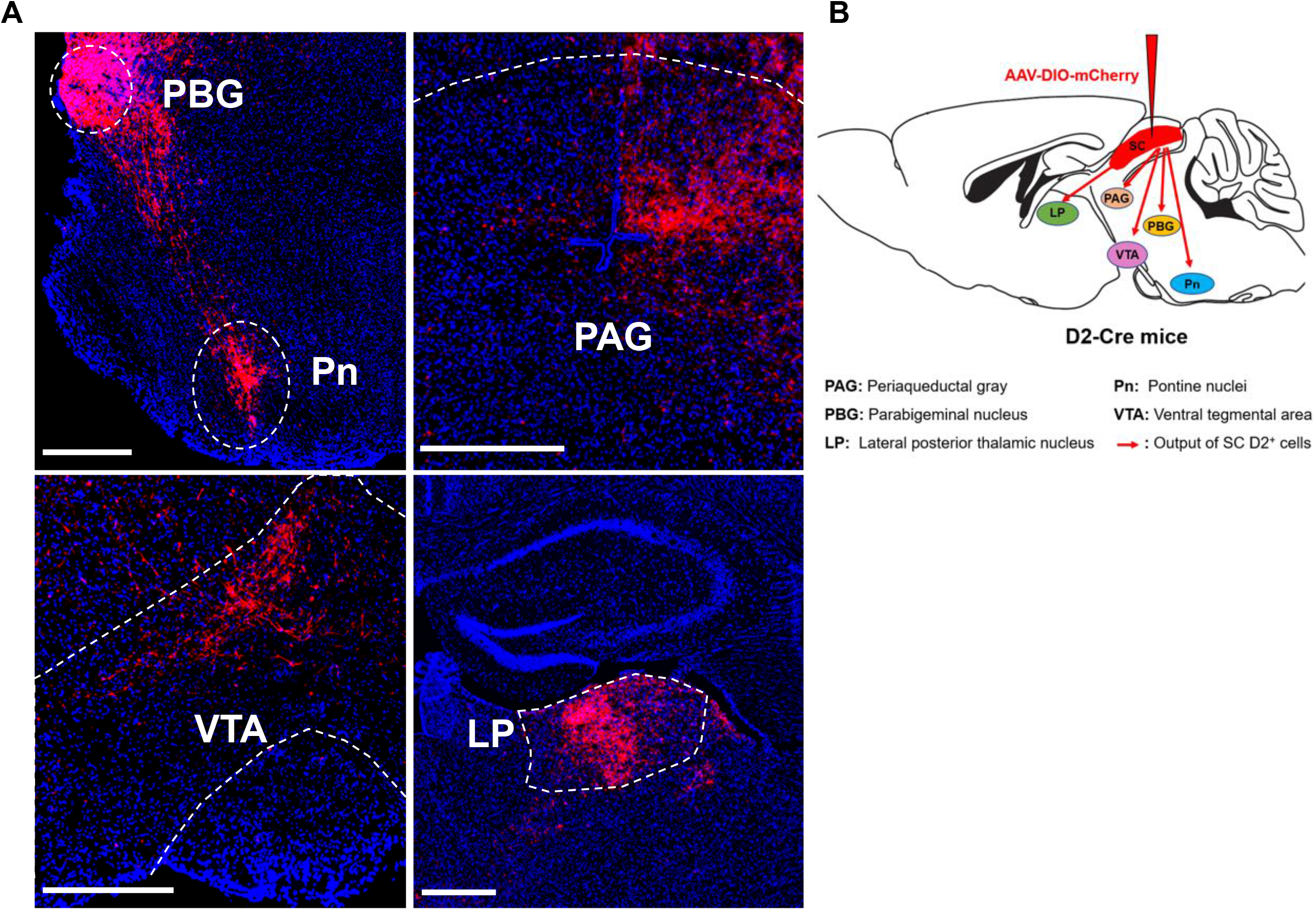
The outputs of SC^D2+^ neuron. **(A)** Anterograde tracing of SC ^D2+^ neurons show fibers in the Parabigeminal nucleus (PBGN), Pontine nuclei (Pn), Periaqueductal gray (PAG), Ventral tegmental area (VTA) and lateral posterior nucleus of the thalamus (LP), and with images of the terminal fibers (scale bars, 500 μm). **(B)** Schematic image of the outputs of the D2-Cre neurons from SC.

**Supplementary Figure 2.**
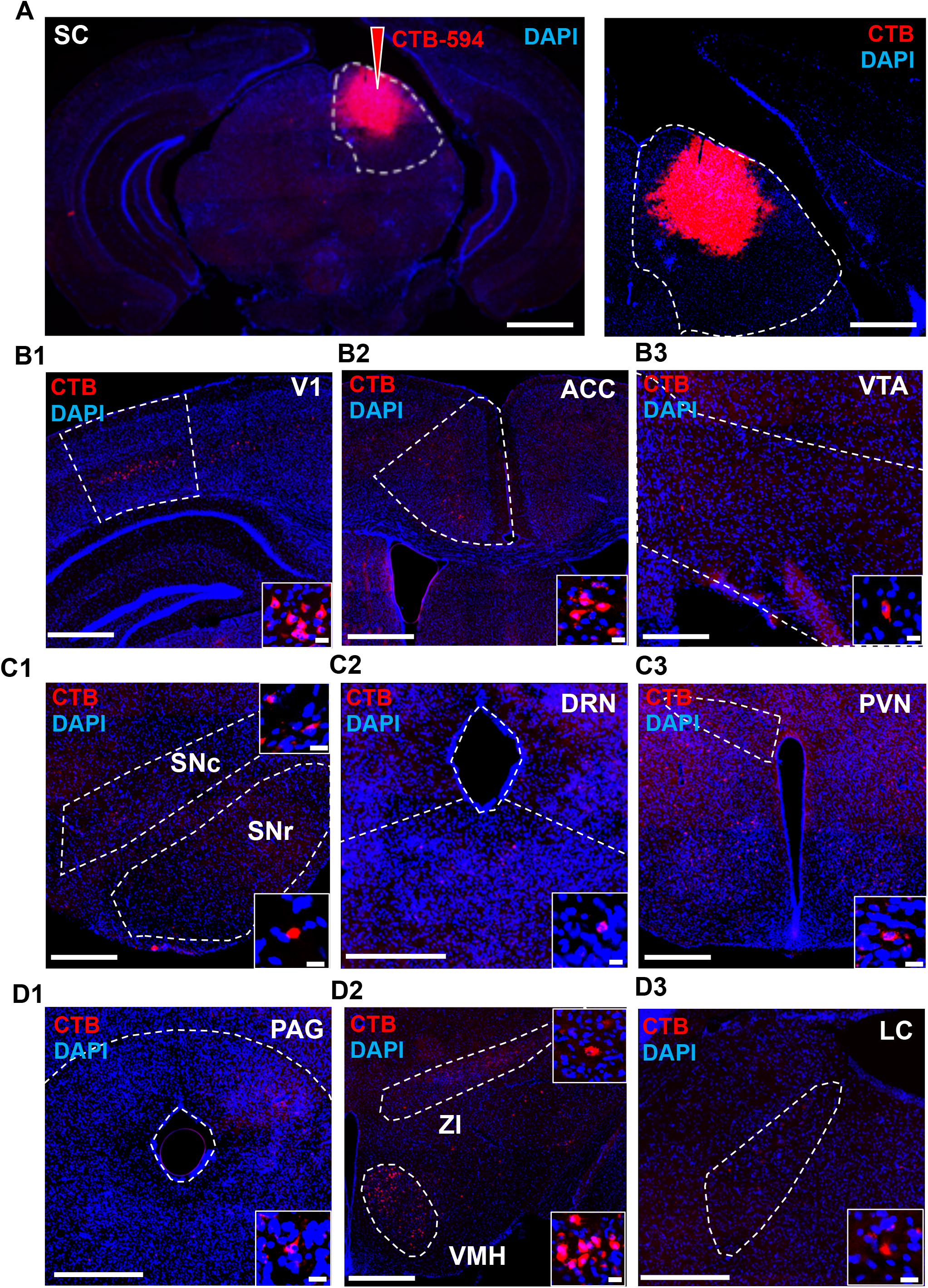
CTB-based retrograde tracing identified the input of SC neurons. **(A)** CTB-594 retrograde tracer was injected into SC. **(B-D)** CTB-594 labeled neurons in regions of primary visual cortex, V1(B1); ACC (B2); VTA(B3); SNc (C1); SNr (C1); DRN (C2); PVN (C3); PAG (D1); VMH (D2); ZI (D2) and LC (D3); (Red, CTB; blue, DAPI, scale bar, 1000 μm, 500 μm and 20 μm, repsectively).

**Supplementary Figure 3.**
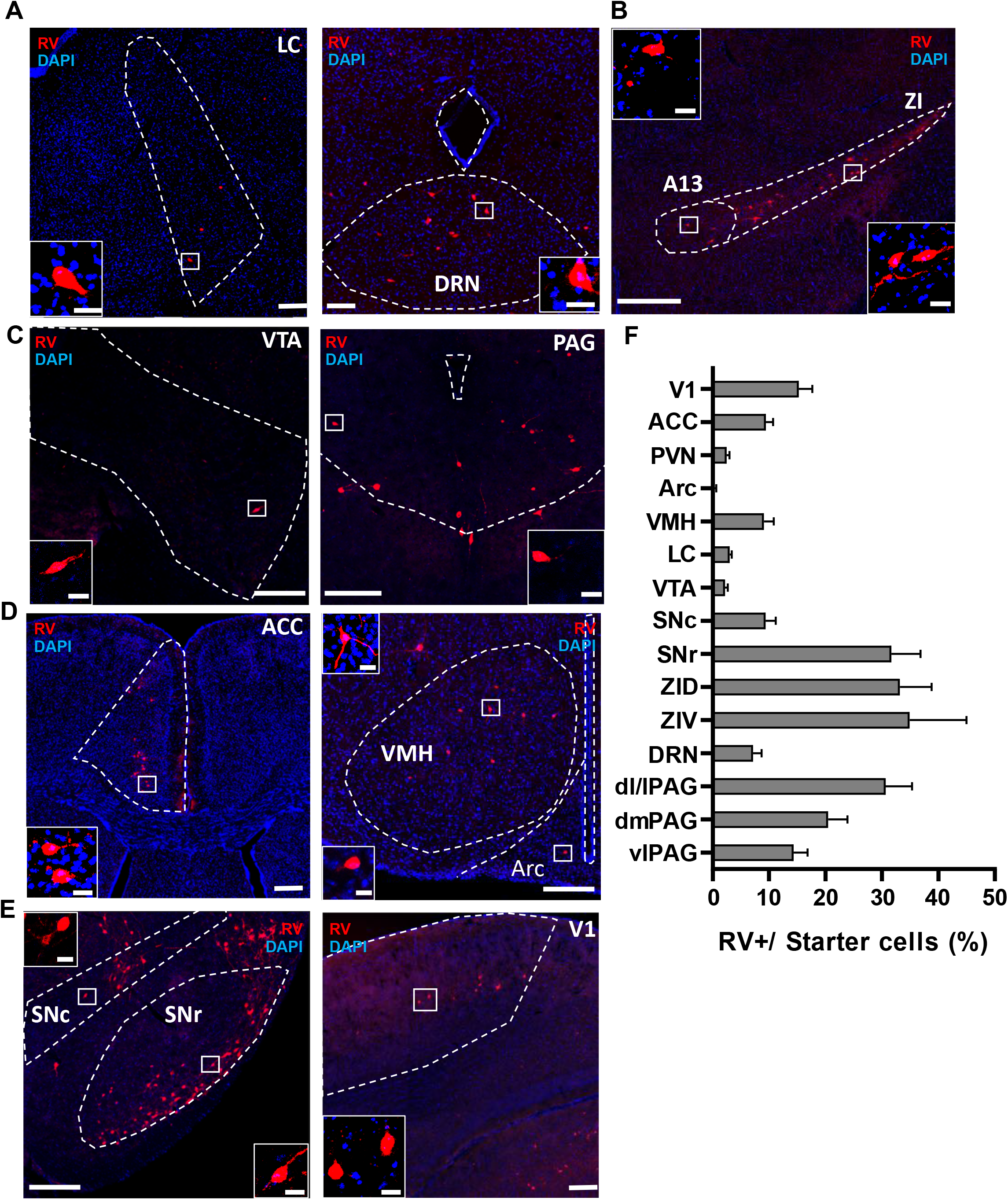
Rabies virus-based viral tracing identified the input of SC ^D2+^ neurons. **(A-E)** Rabies-dsRed labeled neurons in regions of LC, DRN, VTA, PAG, anterior cingulate cortex (ACC), VMH, Arc, SNr, SNc and Primary visual cortex (V1); (Red, rabies-dsRed; blue, DAPI, scale bar, 200 μm and 20 μm). **(F)**Quantification of the number of rabies-dsRed labeled neurons in regions upstream of the SC (n =18-48 slices from 5 mice, data presented as mean ± SEM)

